# Architectural Affordance Impacts Human Sensorimotor Brain Dynamics

**DOI:** 10.1101/2020.10.18.344267

**Authors:** Zakaria Djebbara, Lars Brorson Fich, Klaus Gramann

## Abstract

Action is a medium of collecting sensory information about the environment, which in turn is shaped by architectural affordances. Affordances characterize the fit between the physical structure of the body and capacities for movement and interaction with the environment, thus relying on sensorimotor processes associated with exploring the surroundings. Central to sensorimotor brain dynamics, the attentional mechanisms directing the gating function of sensory signals share neuronal resources with motor-related processes necessary to inferring the external causes of sensory signals. Such a predictive coding approach suggests that sensorimotor dynamics are sensitive to architectural affordances that support or suppress specific kinds of actions for an individual. However, how architectural affordances relate to the attentional mechanisms underlying the gating function for sensory signals remains unknown. Here we demonstrate that event-related desynchronization of alpha-band oscillations in parieto-occipital and medio-temporal regions covary with the architectural affordances. Source-level time-frequency analysis of data recorded in a motor-priming Mobile Brain/Body Imaging experiment revealed strong event-related desynchronization of the alpha band to originate from the posterior cingulate complex and bilateral parahippocampal areas. Our results firstly contribute to the understanding of how the brain resolves architectural affordances relevant to behaviour. Second, our results indicate that the alpha-band originating from the posterior cingulate complex covaries with the architectural affordances before participants interact with the environment. During the interaction, the bilateral parahippocampal areas dynamically reflect the affordable behaviour as perceived through the visual system. We conclude that the sensorimotor dynamics are developed for processing behaviour-relevant features in the designed environment.

## Introduction

When we act, we are changing the perceived environment according to a set of expectations that depend on our body and the environment. The potential ways we can act depends on the affordances of the environment ^1^. Affordances refer to the possibilities for use, intervention, and action which the physical world offers and are determined by the fit between a body’s structure, skills, and capacities for movement and the action-related properties of the environment ^2^. Affordances are thus perceptual and action-related expectations that are systematically reflected in sensorimotor dynamics ^3^. In this sense, cognitive functions that depend on sensory or motor activity are not bound by the physical structure of the body alone, but also by the functional ways in which we interact with the environment.

Preparation and selection of motor action have been studied extensively using frequency-specific oscillatory activity. It is generally held that a vital function of the oscillations in the brain entails transferring information across regions to sustain the binding processes related to various cognitive functions ^4–6^. Particularly, decreases in alpha power relative to a baseline, generally referred to as event-related desynchronization ^ERD, for review:, 7–9^, are consistently reported after a preparatory stimulus in sensorimotor tasks ^10–12^. ERD, as opposed to event-related synchronization (ERS), reflects the release from inhibitory mechanisms and is therefore linked to active cortical processing ^9^. In a traditional motor-related priming task (S1-S2 paradigm), it is held that the processes necessary to interact with the upcoming event occur between the preparatory stimulus (S1) and the imperative stimulus (S2) ^13^. These processes are typically pronounced through ERD between S1 and S2.

Traditionally, the selection of perceived information and the selection of appropriate motor responses has been attributed to attentional mechanisms as separate processes occurring in sequences ^14^. In this sense, attentional mechanisms are responsible for the selection processes regarding sensory information, which then is followed by central executive transformation and selection processes that are then transferred to motor-related mechanisms that implement the selection and execute relevant motor responses. However, recent studies suggest that the preparation and execution of actions and the selection of sensory information unfold in parallel rather than serial-like processes ^3,15–17^. These studies suggest that the selection of motor response and the selection of sensory information may be reflected by the same neural processes ^18^. Therefore, we approach motor attention as sharing the same neuronal resources as visual attention ^19^. The ecological account and predictive coding account of the brain are consistent with such an approach. Accordingly, actions comprise motor predictions based on the sensory consequences of the trajectory of the action ^20,21^, where the shape of these actions depends on the built environment. Therefore, action and perception both shape and are shaped by the environment in ways that comply with a dynamic set of motor predictions, i.e. proprioceptive sensations, and perceptual predictions, i.e. exteroceptive consequences of action ^20,22^.

Cortical oscillations as measured with electrophysiological instruments have for long been associated with such behavioural and cognitive states ^6,7,9,23^. Numerous frequency bands can be dissociated within the power spectrum of the electroencephalogram (EEG), for example, theta, alpha, beta, and gamma—however, their functional significance has caused numerous debates. The emergent alpha rhythm in parietal and occipital areas has been linked to anticipatory attention ^24,25^, re-allocation of resources ^26,27^, inhibition of visual areas to suppress the processing of irrelevant visual information ^28^, top-down inhibition of task-irrelevant brain regions ^6^, and additionally, to the timing of inhibitory processes ^9^. Given the many discovered characteristics of the alpha rhythm ^29^, it may be misguiding to ascribe very general concepts to specific frequency bands ^30^. Instead, our approach lends itself to a biological account of oscillations, where the rhythms are not end-products of a specific cognitive function, but rather components underlying information processing in the brain with the interaction between rhythms revealing more about brain function ^30,31^. Here, we focus on how alpha ERD fits within the processes that involve the selection and control of interactions with the environment. In this sense, we do not cast the cortical mechanisms as semantic processes, but instead as ecological and goal-directed. Thus, the concept of ‘attention’ is here used only relative to the task at hand and cannot be generalized.

We set out to investigate the claim that specific frequencies are modulated in a way that reflects the potential to act in a given environment, i.e. affordances. In a previous study ^3^, we demonstrated this systematic covariation of affordance and brain dynamics in the time domain with modulations of early event-related potentials (ERPs) reflecting the potential to act. Extending on these results, we investigate here how sensorimotor activity is induced relative to motor-selectional mechanisms both (i) upon perceiving the environment and (ii) during the interaction with the environment. By ‘sensorimotor dynamics’ we refer to the brain dynamics when an observer estimates the state of the environment while estimating its own state by integrating sensory and motor information. These dynamics, therefore, reflect one aspect of behaviour. Sensorimotor dynamics comprise changes in the visual, somatosensory, motor, and multisensory areas dependent on aspects of the environment. In this regard, we analysed brain data from an earlier Mobile Brain/Body Imaging (MoBI) ^32–34^ experiment that was designed as a motor priming paradigm ^3^. Participants were located in one of two rooms in a Virtual Reality (VR) environment and were instructed to pass through transitions of varying width connecting the two rooms. A preparatory stimulus (S1) revealed perceptual information about the upcoming transition (whether the transition was too narrow to pass or passable), which was then followed by an imperative stimulus (S2) that revealed whether participants had to actively move through the transition or not. In the interval between S1 and S2, the participants anticipated how to interact with the opening, while S2 instructed participants to pass the transition or remain in the same room. The paradigm thus provides an excellent opportunity to answer the following research question: how are attentional mechanisms relative to the selective processing of actions reflected in the brain? With the aforementioned literature in mind, we expected to find desynchronization over parietal, visual and motor cortices reflecting attention, visual processing and motor activity to covary with the affordances of the environment. We found systematic variations in the parieto-occipital alpha band when participants approached transitions with positive or negative affordance that mirrored the desynchronization patterns. Further examination of the alpha-band activity revealed desynchronization over temporo-occipital areas during perceiving poor affordances (when the transition was too narrow to pass), as compared to the perception of positive affordances (when the transition was passable). These results suggest that the brain dynamics followed a pattern of affordance where the possibility to act determined the type of sensory gating in sensorimotor areas.

## Method

The following description of the experimental paradigm corresponds to the original experiment conducted by Djebbara and colleagues ^3^.

### Participants

The experimental procedure included 20 participants, of which 9 were female (mean age 28.1 years, σ = 6.2 years). All participants were recruited from a participant pool of the Technical University of Berlin, Germany and received either monetary compensation (€10/h) or accredited course hours. None of the participants had a history of neurological pathologies and normal, or corrected to normal, vision with no specific background in architecture. A written informed consent was given and the protocol was approved by the local Ethics Committee and signed by all participants. A single participant was excluded due to technical issues.

### Paradigm

Integrating MoBI with Virtual Reality (VR) allowed for recording and analysing brain activity using EEG in freely behaving human participants moving in virtually designed spaces. The experimentation unfolded in the experimental room of 160 m^2^ in the Berlin Mobile Brain/Body Imaging Laboratories (BeMoBIL). The virtual space was designed to occupy a total of 9 m × 5 m for both virtual rooms, i.e. each room was 4.5 m × 5 m. To complete the task, participants had to transit from one room to a second room—however, doors of different widths manipulated the transition affordances between rooms. The widths of the door in VR varied from impassable (0.2 m, *Narrow*), to passable (1 m, *Mid*) to easily passible (1.5 m, *Wide*). The paradigm was a forewarned *Go/NoGo* paradigm, also known as an S1-S2 paradigm, where S1 was a first stimulus serving as a preparatory signal. S1 presented the participants with the environment including the transition width and S2 was the imperative stimulus. S2 revealed whether participants were allowed to interact with the environment (*Go*), or not (*NoGo*). S2 was pseudorandomized for 50% *Go* and 50% *NoGo*. Therefore, the experimental design was a 3 × 2 repeated measures design where the factors were the type of doors with three levels (*Narrow, Mid, Wide*) and the imperative movement instructions with two levels (*Go/NoGo*). Each participant responded to 240 trials with 40 trials for each factor level. A training phase before starting the 240 trials ensured that participants were comfortable with the protocol and got accustomed to the VR environment. All events in the experiment were registered and collected using LabStreamingLayer ^35^. The main investigator withdrew to the control room and observed the participants through two cameras and a mirrored display of the head-mounted displays the participant was wearing. This ensured minimal-to-none interaction with the participant once the experiment was commenced.

A single trial comprised starting in the dark in the first room inside a predefined starting square directed towards the door (see Figure 1). The participants had to wait for 3 s on average (ITI = 3 s ± 1 s) before the “lights” would go on (S1), so they could perceive the environment including the type of door they had to transit. Facing the closed door, they had to wait for 6 s (ITI = 6 s ± 1 s) before the door would turn green or red for Go or NoGo, respectively. In the case of a Go trial, participants were instructed to walk towards the door, which would slide open when within a distance of 0.3 m, fetch a floating red circle in the second space using their controller, and subsequently return to their starting square. Touching the red circle would elicit a reward of €0.1 added to their reimbursement. If the door was too narrow to pass, they were instructed to try until the walls turned red indicating a collision with the wall resulting in a text informing that they have failed to pass. In the case of a NoGo trial, the participants were instructed to not transition into the second room, but to fill in the emotional questionnaire (Self-Assessment Manikin; SAM) and move on to the next trial. After each trial, they were instructed to fill in the SAM questionnaire irrespective of whether they transitioned through the door or not before moving on to the next trial. The SAM was filled in using a laser pointer from the hand-controller, which also controlled when to turn the “lights off” to move on to the next trial.

**Figure 1.**
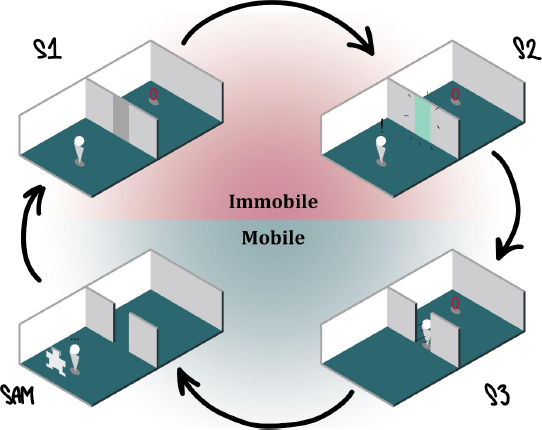
The experiment started as depicted in the top left diagram. Participants were instructed to wait for the lights to come on as they were waiting in the dark (3 s, σ = 1 s). Once the lights came on (S1) they perceived the door they had to pass, however, they were instructed to wait (6 s, σ = 1 s) for the colour of the door to change. Once the colour changed (S2), it turned either green (Go) or red (NoGo). In the case of Go, participants passed the opening (S3), virtually touched the red circle (which in turn releases a monetary bonus), returned to the starting position, and completed the virtual SAM questionnaire. In the case of NoGo, participants turned around and completed the virtual SAM questionnaire. The three different doors had the following dimensions: Narrow, 0.2 m; Mid, 1 m; Wide, 1.5 m. Note the colour codes for each the immobile and mobile phases as they are used throughout the paper.

### EEG Recording

The MoBI approach affords to record human brain dynamics in participants that are actively transitioning the different doors. A mobile 64-channel EEG (eegoSports, ANT Neuro, Enschede, Netherlands) was combined with Windows Mixed Reality goggles (2.89″, 2880 × 1440 resolution, update rate at 90 Hz, 100-degree field of view with a weight of 440 grams) and a high-performance gaming computer back-pack to render the VR environment (Zotac, PC, Partner Limited, Hong Kong, China). The VR environment was scripted in, and powered by, Unity. The participants were equipped with a hand-controller by Acer that was linked to the VR-system (see Figure 2). All EEG data were recorded (DC) with a 0.3 Hz high-pass filter and sampled at 500 Hz with impedances kept below 10 k?. Computational delays were measured by parallel processing a direct event-marker and an event-marker through Unity. The 20 ms ± 4 ms were corrected during the analysis.

**Figure 2.**
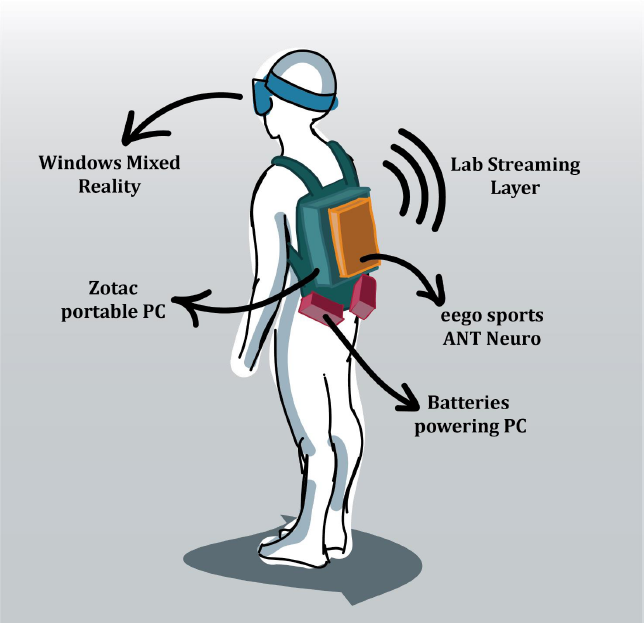
The illustration depicts the MoBI setup used during the experiment. The participants wore a backpack, carrying a high-performance gaming computer (Zotac, blue), powered by two batteries (red). An EEG amplifier (ANT eegoSports, yellow) was attached to the backpack and connected to the computer. The participants wore a VR head-mounted display (Windows Mixed Reality) on top of a 64-channel cap. This setup allowed participants to move freely around while recording data.

### Data analysis

The offline analysis was conducted using MATLAB (MathWorks, Natick, MA, USA) and the EEGLAB Toolbox ^36^. The data were band-pass filtered between 1 Hz and 100 Hz which is within a reasonable range ^37^, and further downsampled to 250 Hz before undergoing automatic cleaning, which consisted of automatically detecting and excluding the most deviant data. Specifically, the data were segmented in epochs of 1000 ms in each dataset where the rejection rate of 18% was based on the mean of epochs and channel heterogeneity. Hereafter, channels with more than five standard deviations from the joint probability of the electrodes were removed and interpolated, then all datasets were re-referenced to an average reference. Adaptive mixture independent component analysis ^38^ was computed on the remaining ranks. This resulted in matrices of ICA spheres and weights. All ICs were associated with an equivalent dipole model as computed by DIPFIT routines ^39^ using a boundary element head model based on the MNI brain (Montreal Neurological Institute, MNI, Montreal, QC, Canada).

Epochs used to analyse the *LightsOn* data epochs were time-locked to the *Lights-On* event (S1) from −500 ms to 4000 ms after the stimulus onset for each factor. Since the imperative stimulus (S2) was presented after 5000 ms, the selected epochs did not contain any brain activity of the S2-onset. Equivalently, epochs used to analyse the *Threshold* (when participants passed the transition) were time-locked to the *Threshold* event (S3) from −4000 ms to 500 ms and thereby describing the brain activity before the event of passing the transition. Approximately 17% of all epochs for the *Lights-On* event while approximately 21% for the *Threshold* event were automatically rejected since they deviated more than five standard deviations from the joint probability and distribution of the activity of all electrodes.

Event-Related Spectral Perturbations (ERSPs) were computed using the *newtimef()* function in EEGLAB. Frequency range from 3 Hz to 100 Hz in log-scale, using wavelet transformation with 2.6 cycles for low frequency and 0.5 cycles for higher frequency. The baseline for the cluster precomputations was defined as −200 ms to 0 ms.

The group-level analysis was computed using all ICs with less than 75% in residual variance of their equivalent dipole model. These ICs, which reflect instantaneous independent time source information, were clustered based on their equivalent dipole locations (weighted = 10), grand-average ERSPs (weight = 5), grand-average ERPs (weight = 1), mean log spectra (weight = 1), and scalp topography (weight = 3) and a region of interest (ROI) located in the occipital cortex. The clustering was driven by a repetitive *k-*means clustering approach (Gramann et al., 2018) with 5000 repetitions to ensure replicability. The number of clusters was determined by ICs per participant so that the total number of ICs (301 ICs) and the total number of participants (19 datasets) yielded 16 clusters. The approach was divided into three steps. First, given our prior results ^3^, the occipital area was defined as ROI with the Talairach coordinates (x = −20, y = −90, z = 7). Second, since each clustering repetition yields a solution, the cluster of interest was selected based on (i) the number of participants with an IC in the cluster, (ii) the ratio of ICs/participant, (iii) the spread (average squared distance) of the cluster centroid, (iv) the mean residual variance of the fitted dipoles, (v) the distance of the x-y-z coordinate of the cluster centroid from the ROI, and (vi) the Mahalanobis distance of the cluster of interest from the median of the total 5000 solutions. These quality measures (i = 4, ii = −3, iii = −1, iv = −2, v =−3, vi = −1) allowed to optimize the clustering solution close to the ROI. Third, the solutions were ranked based on the summed score, where the highest-ranked solution was chosen as the final clustering solution.

Permutation tests (1,000 permutations) using EEGLAB stats were first computed to indicate differences across factor levels. These are visualized in Figure 5 and 6 as the white lines. Based on these results, statistical analysis over group-level ERSPs was computed. These consisted first of averaging the time-frequency data on IC-level for individual participants and then at the cluster level. The time-frequency data were baseline-corrected using the interval of 500 ms before stimulus onset to the timepoint of stimulus onset. Based on the hypothesis, clusters representing parietal, occipital and motor cortices were selected, where clusters with less than 13 participants (75% of total) were excluded. Alpha power values in the frequency band from 8 to 14 Hz were averaged for segments of 500 ms resulting in 8 non-overlapping segments extracted from each total epoch of 4000 ms. For each selected cluster, 3 × 8 factorial ANOVAs were performed testing for power modulation in the alpha frequency band using the door width (3 levels: *Narrow, Mid, Wide*) and the 8 non-overlapping segmented time-windows (8 levels) as the repeated-measures factors. These ANOVAs were computed separately for the motor and parieto-occipital areas for both the immobile and the mobile phase (Figure 4). Significant differences in alpha power were further contrasted by way of Tukey HSD. All ANOVAs were computed as linear mixed models. Contrasts (Tukey HSD) beneath the time-frequency plots in Figure 5 and 6 are based on segmenting the time-frequency data into sets of 500 ms within the frequency range of 8–14 Hz ^40^ for alpha-band and grouping the involved ICs and hereafter take the average. In the case of violations of the sphericity, corrected p-values are reported.

## Results

To address the research question, the two separate experimental phases were analysed, namely a motor-preparatory phase and a motor-execution phase. Given the variation in duration across trials and participants, we epoched the data based on the fastest possible *Go/NoGo*-event. The motor-preparatory phase thus started with the *Lights-On* event and included the first 4000 ms [0 4000 ms] in which participants first saw the room and the transition to the next room. We refer to this event as *LightsOn*. The motor-execution phase was time-locked to the *Threshold* event and included the time period of 4000 ms before crossing the threshold between the two rooms [-4000 0 ms]. We refer to this event as *Threshold*.

### Cluster solution

According to the hypothesis and the condition for inclusion, the clusters of interest included the occipital, motor, and parietal areas, namely *Cluster 3, 6, 9*, and *11* (see Figure 3). Therefore, we selected these clusters for further analysis. In locating the origin of the clusters, the mean of the included ICs was calculated and projected onto the MNI head-model (see Table 1). The cluster representing the cingulate area (*Cluster 3*) was estimated to originate from BA23, which corresponds to the posterior cingulate cortex (PCC). Further, we were able to identify two clusters in each hemisphere in the temporo-occipital region (*Cluster 6* and *11*), which were located to originate from BA19, corresponding to a location near the intersection of the occipital extrastriate areas and the bilateral parahippocampal regions (PHC). A single cluster within the supplementary cortex (SMA) was identified (*Cluster 9*) (see Figure 3). Given the limited spatial resolution of EEG as a neuroimaging method, we interpret the estimated location of the clusters with care. We take them as suggestive rather than absolute. Using the Talairach-coordinates of the mean of each cluster, we labelled the clusters to the nearest grey matter Talairach Client, 41. The involved areas include the PCC, bilateral PHC, and SMA. Since BA23, which corresponds to the retrosplenial cortex (RSC), is within 1 mm of range of the PCC-cluster, it is not possible to exclude the RSC as contributing to the cluster activity—henceforth, retrosplenial *complex* (RSC).

**Table 1.**
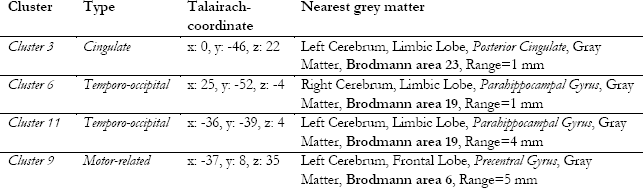
The nearest grey matter of each cluster and their respective Talairach coordinates. The range indicates the distance in millimetres.

| Cluster | Type | Talairach-coordinate | Nearest grey matter |
| --- | --- | --- | --- |
| *Cluster 3* | *Cingulate* | x: 0, y: -46, z: 22 | Left Cerebrum, Limbic Lobe, *Posterior Cingulate*, Gray Matter, **Brodmann area 23**, Range=1 mm |
| *Cluster 6* | *Temporo-occipital* | x: 25, y: -52, z: -4 | Right Cerebrum, Limbic Lobe, *Parahippocampal Gyrus*, Gray Matter, **Brodmann area 19**, Range=1 mm |
| *Cluster 11* | *Temporo-occipital* | x: -36, y: -39, z: 4 | Left Cerebrum, Limbic Lobe, *Parahippocampal Gyrus*, Gray Matter, **Brodmann area 19**, Range=4 mm |
| *Cluster 9* | *Motor-related* | x: -37, y: 8, z: 35 | Left Cerebrum, Frontal Lobe, *Precentral Gyrus*, Gray Matter, **Brodmann area 6**, Range=5 mm |

**Figure 3.**
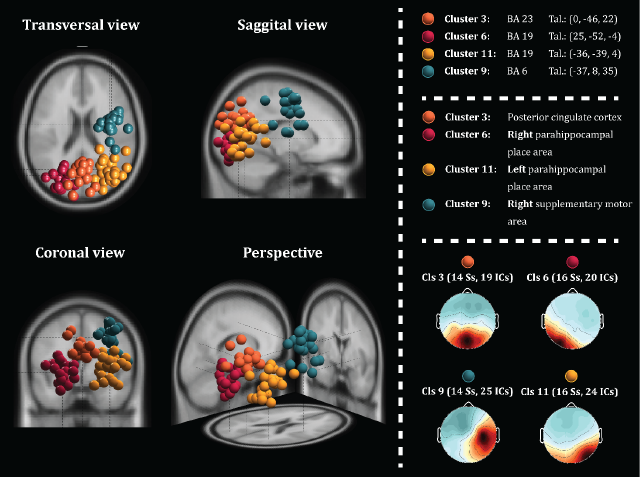
In the left panel, a plot of 3D dipole source locations of Cluster 3, 6, 9 and 11 are depicted onto the MNI model. The centroid of Cluster 3 corresponds to BA23, while both Cluster 6 and 11 corresponds to BA19 in each hemisphere. The centroid of Cluster 9 corresponds to BA6. The lower right panel displays the averaged scalp maps with each its respective number of participants and ICs.

### LightsOn ERSPs

Frequency analyses of the selected clusters during both the *LightsOn* and the *Thresholds* phases revealed clear alpha ERD in the selected clusters (Figure 5). The permutation tests indicated significant differences across conditions in the alpha frequency band. Analysing first the *LightsOn* phase, follow-up two-way repeated-measures 3 (*Narrow, Mid, Wide*) × 8 (time-windows) ANOVAs on alpha-band power for the selected 4 clusters (*Cluster 3, 6, 9, 11*) were computed. *Cluster 3* did not reach significance for the door widths (*F_2,26_ = 2.053, p = .149, μ^2^ = .136*), neither did for the interaction (*F_14,273_ = .775, p = .696, μ^2^ = .038*). Similarly, *Cluster 9* neither reached significant for widths (*F_2,26_ = .36, p = .701, μ^2^ = .027*) nor interactions (*F_14,273_ = .517, p = .923, μ^2^ = .026*). However, *Cluster 6* revealed significant main effects for door widths (*F_2,30_ = 7.202, p = .002, μ^2^ = .324*) without significant interactions (*F_14,315_ = .349, p = .987, μ^2^ = .015*). Equally, *Cluster 11* reached significance for door widths (*F_2,30_ = 3.441, p = .045, μ^2^ = .187*), also without reaching significance for the interaction term (*F_14,315_ = .409, p = .972, μ^2^ = .018*). The posthoc Tukey HSD analyses confirmed that the PHC-clusters (*Cluster 6* and *11*) reflected the affordances of the environment (all p-values are reported in Figure 5). These results suggest that during the immobile period of the experiment, the PHC-clusters reflected the architectural affordances via alpha ERD (see Supplementary Figure 1 for all ERSPs within this phase).

### Threshold ERSPs

Similar for the *Threshold* phase, permutation tests first indicated significant differences across door widths. Two-way 3 × 8 ANOVA repeated measures for each cluster were computed. During this mobile phase, both *Cluster 3* (*F_2,26_ = 4.561, p = .02, μ^2^ = .260*) and *Cluster 9* (*F_2,26_ = 5.021, p = .0143, μ^2^ = .279*) uncovered significant main effects for door width. Interactions for *Cluster 3* (*F_14,273_ = 1.024, p = .43, μ^2^ = .050*) and *Cluster 9* (*F_14,273_ = .22, p = .999, μ^2^ = .011*) did not reach significance. However, *Cluster 6* (*F_2,30_ = 1.028, p = .37, μ^2^ = .064*) and *Cluster 11* (*F_2,30_ = 3.293, p = .051, μ^2^ = .180*) uncovered no significant main effects. The were no significant interactions for neither *Cluster 6* (*F_14,315_ = .753, p = .720, μ^2^ = .032*) nor *Cluster 11* (*F_14,315_ = .736, p = .737, μ^2^ = .032*). Similar to the *LightsOn* phase, the posthoc Tukey HSD analyses revealed that the alpha ERD in the PCC-cluster (*Cluster 3*) and the premotor cluster (*Cluster 9*) reflected the affordances of the environment. The ERSPs of the clusters displayed clear alpha ERD (all p-values are reported in Figure 6). These results suggest that during the mobile session of the experiment, the PCC-cluster and premotor-cluster dynamically reflected the affordances of the environment via alpha ERD (see Supplementary Figure 2 for all ERSPs in this event). For an overview of the relevant clusters in each phase, see Figure 4.

**Figure 4.**
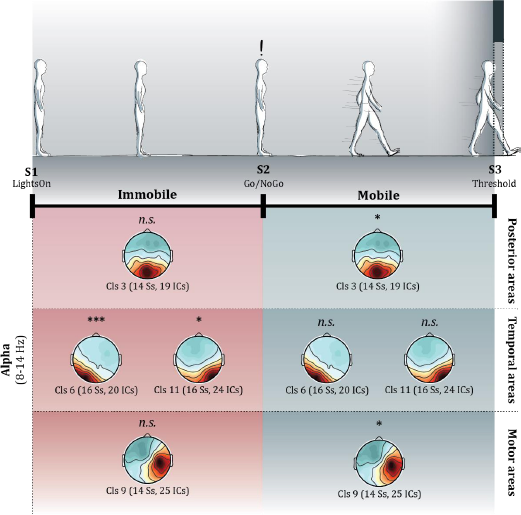
The top panel illustrates the state of the participants while the dense black line represents the timeline. The panel beneath displays the average scalp maps where the asterisk represents the respective ANOVA results (see Results section). The panels are divided by their area to provide an overview of the observed changes.

**Figure 5.**
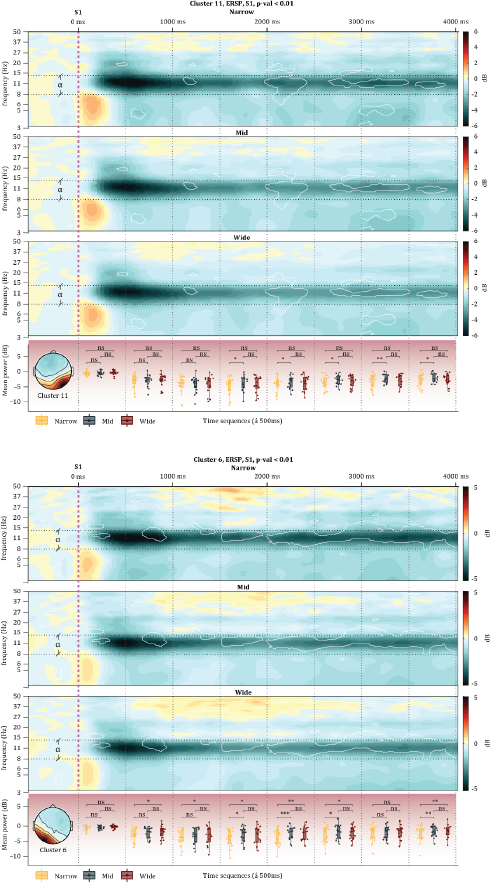
The panel above displays the ERSPs for each condition in Cluster 11 and 6 during the immobile phase. The overlayed white lines represent the areas with a significant difference according to the permutation tests (1,000 permutations). The vertical magenta dashed line represents the onset of the stimulus (Lights On), while the vertical black lines represent the time phases that alpha power values were separated into and analysed by way of ANOVA and Tukey HSD. Horizontal black dashed lines represent the range of the alpha frequency. The bottom panel illustrates the Tukey HSD results; all p > .05 are written as ‘ns’, all p < .05 are written *, all p < .01 are written **, and all p < .001 are written ***.

**Figure 6.**
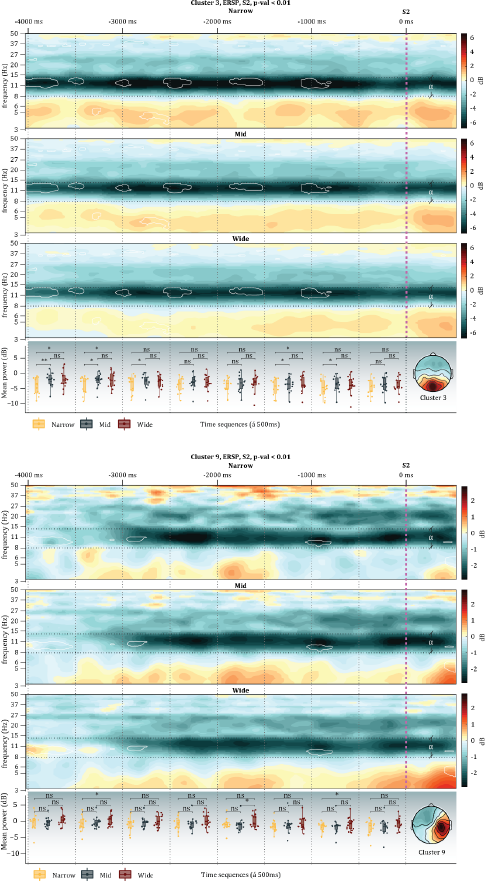
The panel above displays the ERSPs for each condition in Cluster 3 and 9 during the mobile phase. The overlayed white lines represent the areas with a significant difference according to the permutation tests (1,000 permutations). The vertical magenta dashed line represents the moment they passed through the transition (Threshold), while the vertical black lines represent the time phases that alpha power values were separated into and analysed by way of ANOVA and Tukey HSD. Horizontal black dashed lines represent the range of the alpha frequency. The bottom panel illustrates the Tukey HSD results; all p > .05 are written as ‘ns’, all p < .05 are written *, all p < .01 are written **, and all p < .001 are written ***.

## Discussion

With an action-selection approach to attention, we questioned how sensorimotor attentional processes are influenced by architectural affordances. Computing time-frequency analysis and clustering the ICs of EEG data in participants moving through architectural spaces, we found that during the immobile *LightsOn* phase, the bilateral PHC covaried with the affordances of the space—however, while approaching the *Threshold* between the two rooms, we found that the premotor area and PCC reflected the affordances. The deep structures in parieto- and temporo-occipital regions exhibited a significant involvement, which was reflected in the attenuation of the alpha rhythm in all regions. From our clustering solution, we can differentiate between alpha originating from the PCC, bilateral PHC, and premotor cortex. Post-hoc statistical analysis revealed that exclusively the *Narrow* transition was processed differently from the other conditions over the parieto- and temporo-occipital sites. The action-selection process involved with architectural affordances, according to these results, are resolved via parieto- and temporo-occipital alpha that involves a context-sensitive network, which encompasses the PCC, possibly RSC, premotor area and bilateral PHC. We discuss these results in light of the anatomical structures involved in alpha-band attenuation and conclude with a discussion of the behavioural corollary in terms of embodied predictions.

### Behavioural aspect

#### Attention as a balance?

Conservative sensorimotor views cast action as re-actions to stimuli so that action is guided by the stimulus akin behaviourism ^42^. Such an approach favours physical causes above mental causes. However, the ideomotor view stresses goal-directed actions that originate from internal (volatile) causes of action. In this sense, goal-representations, i.e. functional anticipations of actions, play a central role in the emergence of action. Goals and goal-representations are prioritized above stimuli and responses in the generation of action in this view. Nonetheless, in the current study, we conceive action as both dependent on internal and external causes because we take actions to be structures that link movement to goals, and goals to movements ^43^. Such an ecological approach stresses the interaction between the action capabilities of the environment and the physical structure of the body ^1^. It requires parallel internal and external attention as affordances are neither an objective fact nor a subjective feature of our existence. Therefore, since no participant chose to disobey the instructions, this experimental paradigm encompasses a window into the action-selection process (i) upon perceiving the type of transitions (*LightsOn*) and (ii) during the unfolding of the selected action that in turn causes a changing environment (*Go-Threshold*).

In line with the aforementioned critique of ‘attention’ ^30,31,44^, the folk-psychological concept is disregarded, and instead, a biological and phylogenetic understanding is employed. Despite the many studies referring to the parieto-occipital area as the key-node in the attention-network ^45–47^, we stress the role of the parieto- and temporo-occipital area in embodied decision-making processes related to environmental (external) changes ^44,48^. Given an ecological and predictive coding approach, our view of ‘attention’ stems rather from the motor planning process directing the sensory gating function.

#### Embodied predictions

An active inference account of action and perception complies with the ecological foundation of affordances ^49^. In the framework of the free energy principle, the corollary, active inference, cast action and perception as serving the same purpose, namely to minimize our uncertainty about the environment by minimizing free energy ^20^. Central to sensorimotor dynamics, active inference conceives action as the fulfilment of predictions based on inferred states of the ecological environment. The world is present in all its details because the necessary set of actions to bring up a specific perception or information is known and immediately obtainable ^21,50,51^. This essentially means that behaviour can be framed as optimal predictions based on incoming sensory evidence, where the ‘optimal’ is biased by prior preferences or goals. In this sense, embodied predictions ^52^ rests on consequences of sensorimotor contingencies on proprioceptive, interoceptive, and exteroceptive levels where the variable to be controlled resides in the body whereas the controllable states are sensorimotor-related ^53^. By conceiving architectural affordances as embodied predictions, i.e. predictions based on our physical structure, motor signals can be cast under the same neural mechanisms as visual signals. The necessary ‘attention’ in sensory-gating can thus be understood as motor-related attunement where the ‘attention’, i.e. feedback or prediction-errors, reflects a behaviourally relevant selection process from multi-level sensorimotor dynamics. In our case, we observe a strong modulation of the alpha-band as originating from the parieto- and temporo-occipital area.

### Electrophysiological and anatomical aspect

#### Anatomy of thalamocortical alpha

Several studies have shown the visual thalamus to be crucial in the generation and modulation of posterior thalamocortical alpha ^54–61^. Neurophysiologically, the alpha rhythm is viewed as an active inhibitor mechanism gating sensory information relative to perception ^9,24,62^. The thalamus, which holds a large collection of relay neurons, is the only source of information for the neocortex about the body and the environment (safe olfaction). It is connected to the cingulum, which in addition to interconnecting the major regions of the brain also serves as a tract interconnecting with the thalamus, e.g. PCC and retrosplenial complex (RSC). ^63^. It is worth noting that thalamic-cingulate projections posterior to the splenium divide to form separate fascicles in RSC and PHC and further forms a principal route for RSC and PCC projections to the PHC ^64^. However, feedback connections returning to the thalamus, i.e. cortico-thalamic paths, outnumber the outward projections by 5 to 10 fold ^65–67^. Since these inputs relayed through the thalamus arrive via axons with targets often being subcortical motor centres, it raises the possibility that thalamocortical projections also serve as efference copies ^68^. Therefore, the embodied predictions may already be directing the gating function of the thalamus as the sensory information is dynamically collected and processed. This could explain the observed alpha behaviour in the PCC while the participants were approaching the door.

The PCC has been shown to have strong functional connections to many other regions in the brain ^69^, and to be one of the most active regions during rest and during task-related challenges ^70,71^. The cytoarchitectural structure reveals that it is organized to process perceptual input relative to the limbic and hypothalamus regions, which suggests an important role in the internal and external regulations ^72,73^. Accordingly, the PCC has been ascribed to the capability of controlling the balance between the internal ‘attention’, e.g. recalling autobiographical memories, and external ‘attention’, e.g. environmental changes. Particularly the PCC and RSC have been suggested as key areas in this balance ^74,75^. Given that studies using nonhuman primates have demonstrated the importance and responsiveness of the PCC in environmental changes ^48^ and the alterations of behaviour ^76^, the balancing of internal and external attention may be a plausible explanation for the involvement of the PCC in this study. This is consistent with the ‘affordance competition hypothesis’, which suggests that as the environment changes, the affordances change along, which is then manifested in dynamic embodied predictions that propel the body effortlessly through the environment ^77,78^.

However, with the low spatial resolution of the EEG, we speculate whether the origin could be in an adjacent and closely related area. For instance, Bonner and Epstein ^79^, in a functional neuroimaging study, found that a scene-selective region of the visual cortex, labelled the occipital place area (OPA), could be used to predict the affordances of scenes. Studies in nonhuman primates propose that the nature of the connectivity of the cortex in the middle to dorsal levels of the parieto-occipital sulcus, specifically the V6/V6A areas, to be involved in visual guidance of movement ^80,81^. The involvement of the visuomotor area V6 has further shown to receive its primary afferent from thalamic nuclei providing visual and somatic inputs ^82^ and further suggested to be heavily engaged in sensorimotor integration ^80^. As the PCC-cluster is exclusively observed to covary with the affordances during movement, the V6/OPA area cannot be excluded. Therefore, the V6/OPA also qualifies as a source for the observed variation—nonetheless, both the V6/OPA and PCC are theorized to interact directly with the thalamus.**Modulation of PCC alpha.** We suspect the observed posterior alpha during *LightsOn* in the PCC-cluster to be of cortico-thalamic feedback origin. Interestingly, the PCC-cluster showed significant differences across affordances only while performing the task of acting in the environment (*Threshold*), which is arguably ideomotor-dominant, as opposed to the anticipative interval (*LightsOn*) that is sensorimotor-dominant. We speculate whether this difference in the type of task reflected in the PCC-alpha reveals the nature of affordances as the balancing between internal and external ‘attention’. Altogether, with the central role of the thalamus in sensorimotor integration and the PCC in the balancing of internal and external ‘attention’ in mind, we propose that the alpha-band attenuations can be interpreted as an indirect measure of sensorimotor demand, i.e. the demand of neuronal processing for estimating the environment and own state by integrating sensory and motor information. The motor information may then be adjusted by the cortico-thalamic embodied predictions in the thalamus and thereby guiding the gating function ^68^. This means that the subsequent cortical response is modulated already at the entry to the cerebral cortex. Indeed, if the source of the alpha rhythm during movement originates from the PCC, we would expect to find alpha ERD to covary with the sensorimotor demand. For that reason, the posterior alpha serves as an excellent marker of dynamic action-selection processes relative to the affordances as actions unfold.

#### Modulation of premotor alpha

Although we observe ERD in the alpha-band over premotor cortex after the *LightsOn* event, the differences during this phase did not reach significance. The presence of clear alpha ERD suggests activity over the premotor cortex, however, this activity is not modulated by the affordances of the transition but suggests rather that one is in general preparation. Instead, during the *Threshold* phase, the activity seems to reflect the affordances of the door suggesting that the premotor cortex is highly involved in the continuous thalamic action-perception feedback loop suggested above. A previous study shows stronger alpha desynchronization as emerging from BA6 during visually guided reaching, which is equally observed in our recordings ^10^. In summary, the activity of the premotor cortex is not modulated by the affordances during preparation but rather involved during the active phase.

#### Modulation of PHC alpha

Visuospatial processing in the PHC area has in numerous functional neuroimaging studies demonstrated selective activity when viewing scenes or environments, which has linked it to our ability to recognize and navigate the world ^79,83–85^. Particularly the perception of buildings and large-scale spaces has already been demonstrated to be linked with the activity of the PHC area ^85–87^. A recent review suggests a context processing network involving a dynamic relation between the PCC, RSC, PHC and OPA ^88^. Interestingly, we identified the same areas—however, our interpretation stresses the role of the thalamocortical-corticothalamic interactions, particularly due to the recorded alpha-band ERD. Crucially, we observe condition-specific sustained alpha ERD exclusively during the passive *LightsOn* event in the PHC areas. In line with the existing theories, we interpret the activity as the processing of contextual information—however, we suggest that since the available sensory information provided by the thalamus is affordance-sensitive, the PHC activity reflects the associated behaviours as possible in the environment rather than place- or location-specific information processing. Since the contextual information is continuously and correctly predicted as the participants start acting, the PHC area shows no condition-specific differences as opposed to the PCC. According to the thalamic projections described above, we interpret the observed alpha-band activity in PHC to stem from thalamic interactions through the cingulum and the thalamus-PCC-PHC link.

## Conclusion

In the current study, it was asked how architectural affordances relate to the attentional mechanisms underlying the gating function for sensory signals both upon perceiving the environment and during the interaction with the environment, are induced. Assuming that the magnitude of the alpha oscillations reflects the impact on information processing in the brain, the results suggest the parieto- and temporo-occipital alpha ERD serves as a marker for sensorimotor integration during interaction with the environment. For instance, while approaching staircases, passing through doors, and turning corners are all examples of situations that require dynamic processing of architectural affordances. Once the corner is turned, the processing of the new environment, our results suggest, is reflected in the medio-temporal alpha ERD. We suggest the PHC area to be involved in the immediate processing of affordances. Particularly the interaction between the PCC, possibly the RSC, and the bilateral PHCs via cortico-thalamic alpha ERD suggests an action-perception mechanism sensitive to the architectural affordances. Interestingly, it is the same pattern that emerges here as in the study by Djebbara et al. ^3^. Additionally, by conceiving the PCC as a transmodal thalamocortical hub, the PCC becomes crucial to the balance between internal and external ‘attention’, i.e. the breadth of internal and external selection of actions. Therefore, we interpret the visual sensory processing as inherently biased by an understanding of the body by internal attention and the environment by external attention.

Indeed, moving in space is to continuously construct a prediction of a world that we perceive as dependent on our action potentials that in turn is manifested in cortical oscillations. This suggests that users of space hold a principle of anticipation that architects should keep in mind when designing the context for actions. In this sense, by designing our environments, architects design cortical activity. However, we still have a long road ahead to decently understand how architectural affordances impact various levels of brain dynamics.

### Further research

Provided the thalamocortical origin in both the parieto-occipital and temporal regions, we speculate how architectural design may implicitly, but constantly, influence subcortical structures that in turn project to numerous other regions in the brain. A critical factor in the analysed data is the brain activity of behaving human beings. Indeed, using architectural design as a medium for investigating action-selection and sensorimotor integration is an exceptional approach, which calls upon more experimentation. Here, the MoBI-approach proves an excellent way forward. Assisting such experimentations, particularly in articulating the type of involvement of the known subcortical networks, we propose the use of dynamic causal modelling ^89^ to allow for multiple models to compete and infer which of the models best explain the acquired data.

## Acknowledgements

We would like to thank the Berlin Mobile Brain/Body Imaging group, including Marius Klug, Lukas Gehrke, Anna Wunderlich and Federica Nenna for their technical assistance and many fruitful discussions.

---

**Supplementary Figure 1.**
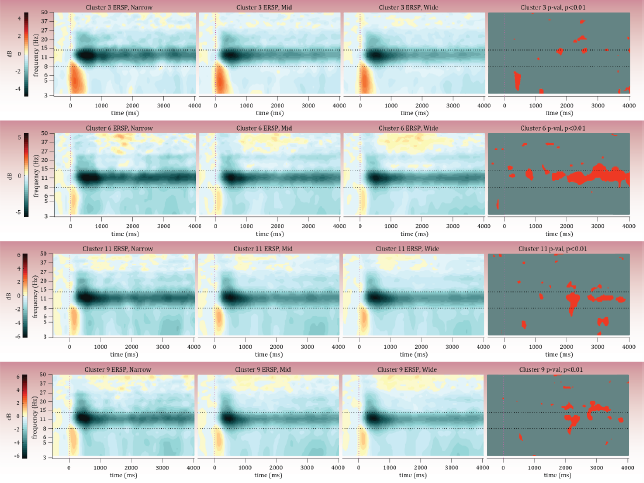

**Supplementary Figure 2.**
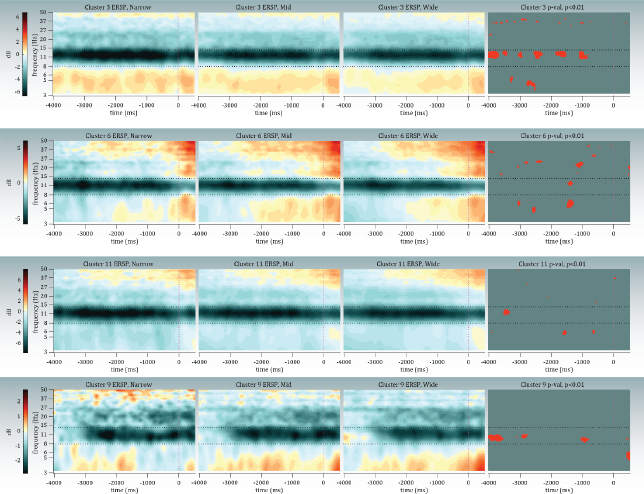

